# Intracellular softening and fluidification reveals a mechanical switch of cytoskeletal material contributions during division

**DOI:** 10.1101/2021.01.07.425761

**Authors:** Sebastian Hurst, Bart E. Vos, Timo Betz

## Abstract

The life and death of an organism depends largely on correct cell division. While the overall biochemical signaling and morphological processes during mitosis are well understood, the importance of mechanical forces and material properties is only just starting to be discovered. Recent studies of global cell stiffening during cell division may imply an understanding of the cytosol mechanics that is mistaken. Here we show that in contrast to the stiffening process in the cell cortex, the interior of the cell undergoes a softening and fluidification that is accompanied by a decrease of active forces driving particle mobility. Using optical tweezers-based microrheology we capture the complex active and passive material state of the cytoplasm using only six relevant parameters. We demonstrate that the softening occurs because of a surprising role switch between microtubules and actin, where the intracellular, actin-based mechanics is largely controlled by a formin-mediated network.

## Introduction

Cell division is a critical process during the cell cycle marked by drastic and well-controlled changes in the cells’ biochemistry, morphology and material properties. Slight deviations during this fragile phase can result in severe cellular damage and can lead to cell death or mutations that encourage the formation of cancerous cells [1]. During division, most cells undergo tremendous morphological changes [2, 3], such as rounding into a sphere-like shape to ensure correct spindle formation [4], with few known exceptions [5]. As morphological changes are driven by a drastic remodeling of the cytoskeleton [6, 7, 8] and stiffening of the cell cortex, a naive view leads to the conclusion that this stiffening occurs throughout the whole cell. This notion has been supported by external mechanical probe experiments that show a critical pushing force and stiffening of the cortex, which is thought to prepare the intracellular space for correct division [9, 10, 11]. Typically, this is referred as mitotic cell stiffening, which implies stiffening of the whole cell, including the interior of the cell. However, these important insights from external AFM measurements become misleading when they are applied to the properties of the cytoplasm inside the cell. As the thick and rigid actin cortex prevents experimental access, intracellular mechanics are not measurable with external probes. Despite this limited accessibility, for a proper understanding of the mechanical processes during division, detailed knowledge of the intracellular viscoelastic material properties is indispensable, as these properties can have a great impact and even indirectly regulate cell fate [12, 13].

Based on the morphological changes and the organizational restructuring, the intracellular mechanical properties would be expected to change during cell division. Further, the striking cortical changes demonstrate that during division the cells’ physical parameters are well controlled, hinting for a crucial role for mechanics in division. Indeed, in this study, we show that changes to the intracellular mechanics during mitosis do occur, as we measured both, viscoelasticity and active, non-thermal transport. To access the intracellular properties felt by endogenous objects, we probed intracellular particles using optical tweezers-based microrheology. This precise method allows for applying an oscillating force without harmful interference, while precisely measuring the resulting particle motion. The direct experimental readout is the complex shear modulus *G*\*(*f*), which fully quantifies the dynamic viscoelastic material properties and is represented as a frequency dependent quantity [14, 15, 16]. Using phenomenological models, the viscoelastic shear modulus can be interpreted using only a few relevant parameters, which allows separating out the different contributions to the mechanical changes that occur during mitosis.

In addition to the passive mechanics, cells as living objects are active systems and constantly generate forces, especially during mitosis [17, 18]. The resulting non-equilibrium situation complicates the physical description, as it provides an active component to particle and organelle mobility that acts in addition to classical Brownian diffusion and often dominates the transport properties. While the strength of Brownian motion can be directly predicted from the viscoelastic material properties, the active transport is under the control of the cell. However, these active transport processes can be directly quantified by comparing the measured particle motion with the expected Brownian diffusion [19, 20, 21]. Disentangling passive and active diffusion is highly relevant, as only the active transport can be regulated by the cell. A simple, yet useful interpretation of the combined effects of Brownian and active transport forces is quantifying the mechanical energy required to explain the observed motion taking into account the measured material properties.

In this study, we show that both mechanics and intracellular activity change drastically in dividing epithelial cells throughout mitosis. The cytoplasm becomes softer, with an increased fluidity, and is less active going into metaphase, while it stiffens and increases its effective mobility again towards the end of division. Hence, the mechanical properties of the cell cortex and the cell interior show an inverse behaviour. Surprisingly, a detailed analysis reveals that between interphase and mitosis actin and microtubules switch their roles in cytoplasm mechanics.

## Results and discussion

### Cytoplasmic mechanics in mitotic cells follow the generalized Kelvin-Voigt model

The fundamental reorganization of the cell during mitosis and the known cortical stiffening suggest that the intracellular material properties also adapt to support mitosis. To address this question, we performed intracellular active microrheology on dividing epithelial MDCK cells throughout the different phases of mitosis (Fig. 1a,f). As probe particles for optical tweezers measurements, we used 1 μm sized phagocytosed particles; we used these particles because they are membrane-encapsulated and, hence, provide a good proxy for the intracellular mechanics experienced by organelles and vesicles, which are typically also membrane-bound.

**Figure 1:**
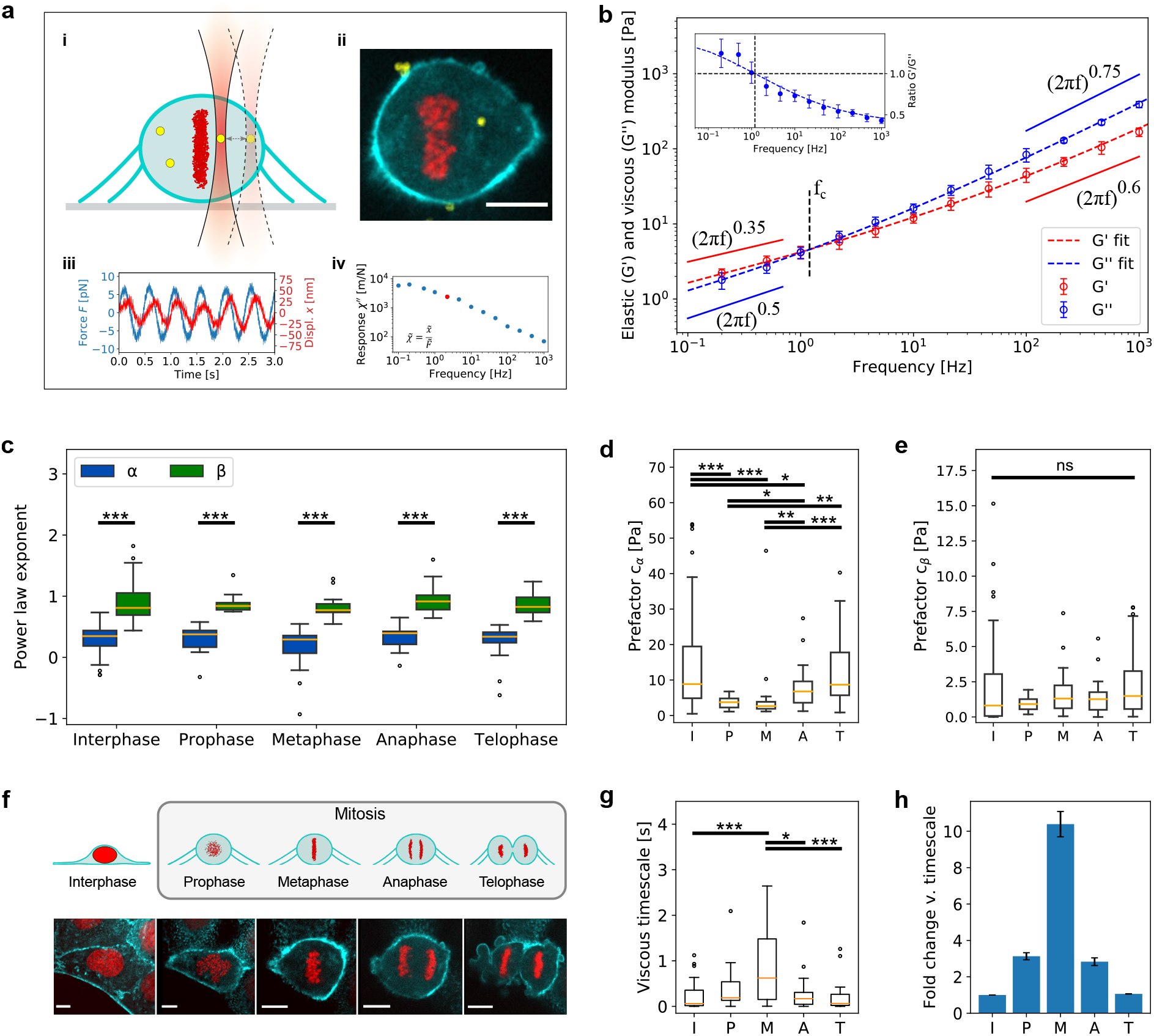
Intracellular viscoelasticity in mitosis. Active microrheology measurements and fractional viscoelastic fit parameters describing the mechanics in mitosis. **a)** Microrheology on phagocytosed particles. i) schematic of dividing cell with particle. ii) dividing MDCK cell expressing H2B-mCherry (red) and Lifeact-GFP (cyan) with phagocytosed particle (yellow). iii) representative measurement at 2 Hz with oscillating applied force and resulting particle displacement. iv) response at 13 frequencies ranging from 0.1 Hz to 1000 Hz. Measurement from iii) shown in red. **b)** Elastic and viscous modulus with fit of fractional viscoelastic model to data of cell in metaphase. Inset: ratio of G’ and G”. **c)** Power law exponents *α* and *β* of viscoelastic fit for different phases of mitosis. They are significantly different in each phase, but *α* and *β* remain constant throughout mitosis. **d)** Statistics of the prefactor *c_α_* of viscoelastic fit shows that the solid contribution is drastically reduced during pro- and metaphase and recovers the interphase value towards telophase. **e)** In contrast, the prefactor *c_β_* of viscoelastic fit shows no significant changes during the whole process. **f)** Top: schematic of dividing cell illustrates the observed process (side view), bottom: fluorescent images with H2B-mCherry (red) and Lifeact-GFP (cyan) demonstrate the drastic organizational changes (top view). Cells were categorized according to H2B-signal. **g)** Detailed analysis of the viscous timescale *τ* = 1/f_*c*_, below which the material is dominated by viscous processes, shows a significant increase. **h)** The relative increase of the viscous timescale *τ* shows that fluid-like dissipation extends to much longer timescales during metaphase as compared to interphase. Scale bars: 10 μm. n_*cells*_ = 58, 15, 19, 21, 20.

The result of active microrheology yields the complex response function *χ*\*(*f*) (Fig. 1a), which can be used to determine the frequency dependent complex shear modulus *G*\*(*f*) via the generalized Stokes-Einstein relation [19]. Active microrheology is required, as passive approaches neglect the active force generation inside the cells. However, the resulting complex moduli (Fig. 1b), are difficult to interpret and compare. Therefore, we model the real (G’) and imaginary part (G”) of the shear modulus in terms of a fractional two element approach using a generalized Kelvin-Voigt (GKV) model [22]: *G*\*(*f*) = *c_α_* · (*i*2*πf*)^*α*^ + *c_β_* · (*i*2*πf*)^*β*^ (Fig. 1b). Here, we extend previously used single power law approaches [23, 24, 25, 26, 27] and remain consistent with recent approaches to describe cellular mechanics [28, 29, 30, 31]. The big advantage of such a model is that it reduces the complex frequency dependent measurements to four parameters, giving a full, physically consistent description of the complex shear modulus. Besides providing an excellent fit to the shear moduli, with mean 〈*r*^2^〉-values of 0.97, the model also nicely describes the ratio of G’ and G” (Fig. 1b, inset), thus illustrating whether the elastic or the viscous contributions dominate the overall mechanical properties. Our results show that fast, high-frequency motion behaves more viscously, while at low frequencies the elastic contribution dominates. This is directly explained by the notion that fast motion provides more dissipative friction. The switch from elastic to dissipative behaviour is easily quantified by the crossover frequency *f_c_*, which marks a viscous timescale *τ* = 1/*f_c_*. Hence, for any motion on timescales smaller than *τ* the viscous dissipation dominates.

The GKV model enables a physical interpretation of the extracted parameters in terms of two different and independent materials. The already introduced distinction between elastic low-frequency and dissipative high-frequency behaviour is also instrumental here, as it implies a solid-like material defined by *c_α_* and *α*, and a fluid-like material defined by *c_β_* and *β*. A similar model was introduced as a poroelastic interpretation of the cytoplasm [32]. This is consistent with the results indicating that in the low-frequency region the elastic modulus dominates, while in the high-frequency region the viscous components become prominent. Although the nature of each contributor remains undefined in the model, the prefactors *c_α_* and *c_β_* and the power law exponents *α* and *β* provide direct insight into the material’s characteristics. Additionally, the model hypothesis of two interpenetrating materials directly suggests the testable hypothesis that the solid and liquid components can be modified independently of each other.

Although the mechanical properties inside mitotic cells had never been addressed until now, our results (Fig. 1) are consistent with a series of previous measurements on non-dividing cells [30] and might be interpreted as the result of classical semiflexible filament mechanics which predicts a power law exponent of 0.75 for the fluid component [33, 34]. This is a fashionable view, as actin filaments are thought to be the main mechanical components inside cells. However, this understanding is challenged by our and other recent results. Although the here observed power law exponents are close to 0.75, they still remain larger in interphase and mitosis (Fig. 1c). Additionally, removing the f-actin in mouse oocytes shows that the mechanical properties of the cytosol are independent of actin [16]; we also confirmed this in interphase cells (Fig. 3d). This questions the interpretation that such an exponent can be directly attributed to the actin cytoskeleton in the cytosol, a point we address later in this study. Overall, these results demonstrate that the mechanical properties inside mitotic cells can be modeled by a two-viscoelastic material approach.

### Active microrheology reveals intracellular softening and fluidification during mitosis

As the cell cortex stiffens during mitosis [9, 11], we wondered if the same also applies to the inside of the cell. To test this, we performed the described active microrheology analysis in interphase and during ongoing mitosis. Recording both the actin cytoskeleton using Lifeact-GFP and the DNA using H2B-mCherry during the measurements allowed us to precisely attribute the measurements to the different phases (Fig. 1f). First, our data confirmed that the introduced GKV model provides an excellent description of the cytosol mechanics throughout mitosis and that we continuously see two characteristic contributions of a solid- and a fluid-like material. We find not only that for each phase the exponents *α* and *β* are significantly different, but also that each of them remains surprisingly constant throughout mitosis (Fig. 1c). This is unexpected, given both the drastic change of cell shape and the massive reorganization of the cytosol. As the power law exponent represents a fingerprint of the material characteristics, the observed stability suggests that either these parameters are highly regulated, or that the high- and low-frequency contributions are so different that the restructuring of the cell during mitosis does not change their respective natures.

Next we focused on the evolution of the prefactors during mitosis. The known stiffening of the cortex has been often interpreted as whole-cell stiffening, which implies that the cytoplasm will also stiffen. This would predict an increase of *c_α_*. To our surprise, we saw a strong, and highly significant decrease in the contribution of the solid-like material to the overall viscoelastic material properties, manifested in a reduction of *c_α_* (Fig. 1d). The corresponding value collapses from *c_α_* = (8.84 ± 1.75) Pa in interphase (n = 58) to *c_α_* = (2.64 ± 2.26) Pa in metaphase (n = 19), which is a reduction to less than 30 % of the interphase value (see also Supplementary table S1). In contrast, the prefactor of the fluid-like material remained unchanged during mitosis with a median value of *c_β_* = (1.11 ± 0.22) Pa (n = 131, Fig. 1e) throughout all phases. This directly shows that the low-frequency stiffness is drastically reduced, and as a consequence the relative contribution of the fluid-like material becomes more dominant. As a result, the viscous timescale increases during mitosis (Fig. 1g), suggesting that the fluid characteristics become more important as they are dominant for longer timescales. This can be interpreted as a fluidification of the cytoplasm, which is highlighted by a more than tenfold relative increase of this timescale when compared to interphase (Fig. 1h). Towards the end of mitosis, namely at telophase, all these differences vanish, and the extracted material parameters show no significant difference to interphase. To validate that these results are not MDCK specific, the measurements were repeated in HeLa cells (Supplementary fig. S1). We found the same overall behaviour regarding softening and fluidification during mitosis, although the absolute numbers of prefactors and exponents varied.

The robustness of the results suggests that the reduced contribution of the solid-like material and the resulting fluidification is a general phenomenon that is either functionally relevant or a byproduct of the structural changes inside the cell during mitosis. Functionally, these changes can mechanically simplify the correct targeting of the microtubules to the kinetochores and, moreover, the separation of the chromosomes. Furthermore, the intracellular softening gives an additional functional role for the cortical stiffening, which protects the now softer inside of the cell from external mechanical perturbations. An interesting observation is that the mechanical properties already recover to their interphase values before mitosis has finished.

### Intracellular activity decreases in mitosis

The pure mechanical effect of the observed fluidification helps to reduce the cellular forces required to position chromosomes and cell organelles in mitosis. This suggests that the effective energy spent by the cells in transport processes is greatly reduced throughout the mitotic reorganization. To test this hypothesis, we directly measured the mechanical energy required for organelle-like particle mobility and compared this to the thermal energy available for Brownian diffusion. This active energy can be determined by combining the material properties with the observable particle motion. Spontaneous particle motion in the absence of the trapping laser was measured with nanometer precision using back focal plane detection by a low-power, non-trapping detection laser (Fig. 2a, inset). From the particle motion, the free power spectral density 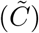 was calculated (Fig. 2a). By comparing the free motion 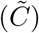 with the expected Brownian motion, the active energy *E_a_*(*f*) can be accessed in units of thermal energy as 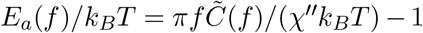. Plotting the rescaled spontaneous fluctuations and the response function together directly shows that in the high-frequency range, the particle mobility can be explained by thermal motion, as both graphs overlap (Fig. 2b). However, in the low-frequency regime, active processes drive an increase in particle motion (shaded area in Fig. 2b). This enhanced motion is also direct proof that the fluctuation-dissipation-theorem (FDT) is violated in the low-frequency regime, which is consistent with a number of recent measurements in interphase cells [35, 36, 31, 16], and confirms that passive microrheology approaches are not valid in mitotic cells. The violation of the FDT in the low-frequency regime is attributed to activity generated by the cell metabolism and transport processes, which we here refer to as ‘active motion’. Consequently, the active energy is a direct measure of this active motion. Inspection of the active energy shows that it follows a power law dependence, hence allowing us to describe it with a power law of the form *E_a_*(*f*) = *e*_0_*f ^ν^* for every mitotic phase (Fig. 2c). Reminiscent of the process for the mechanical properties, we obtain two parameters that quantify the active energy, where a prefactor *e*_0_ gives the strength of the active, metabolic contribution to the motion, while the power law exponent *ν* describes the nature of the active mobile forces acting on the particles. This result allows us to fully describe the mechanical state of a cell, including its activity, with only six parameters.

**Figure 2:**
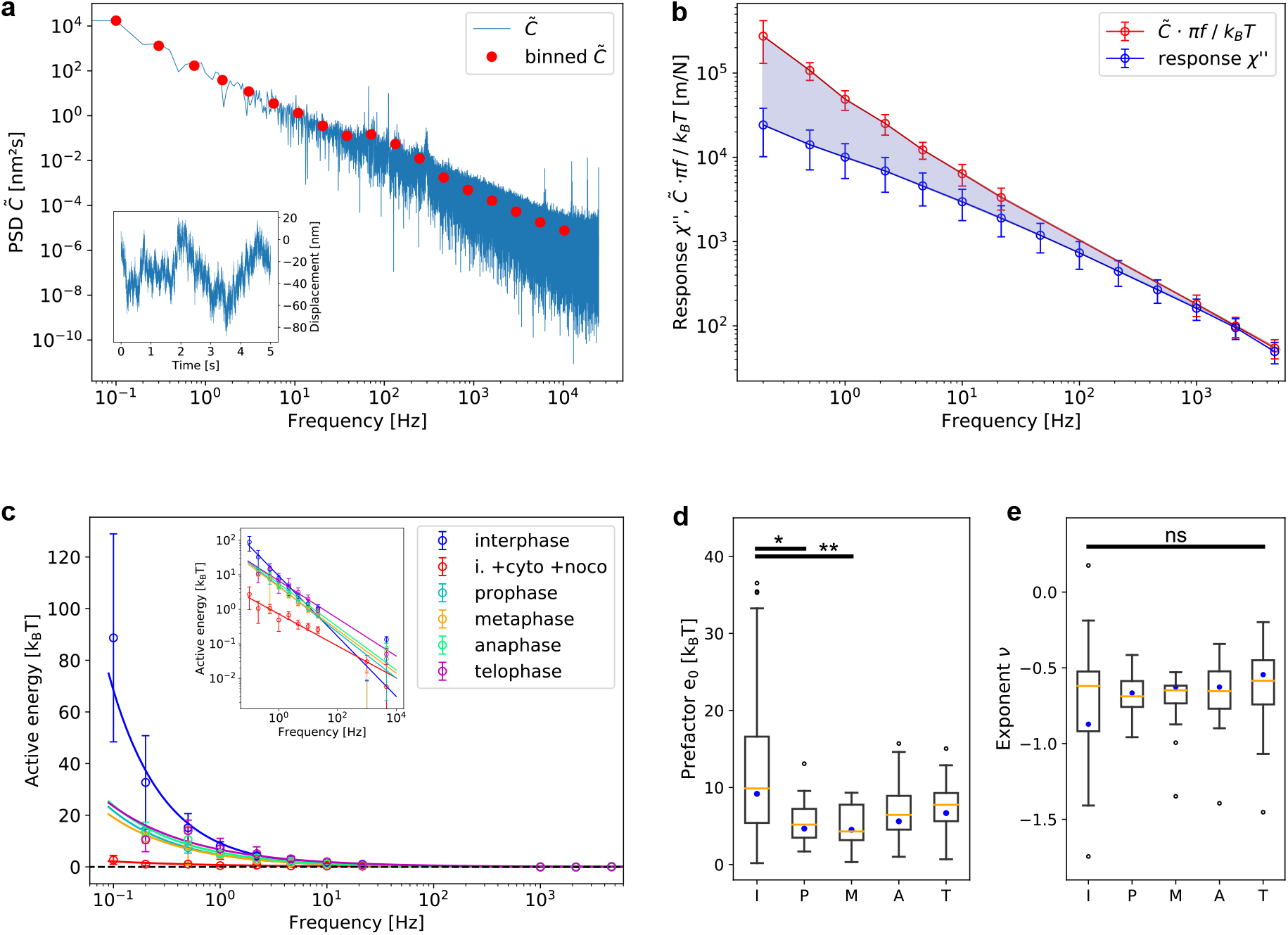
Spontaneous fluctuations and active energy in dividing cells. Responses from active microrheology and spontaneous fluctuation analysis are combined to get information about the intracellular activity. **a)** The power spectral density 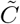 quantifies the spontaneous motion of the particles. We show the un-binned, hence, raw data and the data as binned to the frequencies relevant for comparing them with active microrheology measurements. Inset: Time-dependent example track in one direction of a spontaneously fluctuating particle. **b)** The combination of active microrheology and spontaneous fluctuations demonstrates the effect of active processes driving motion in the shaded area. The violation of the fluctuation-dissipation-theorem indicates the active contribution from the cell by metabolic activity. **c)** Active energy of different mitotic phases calculated from quotient of active microrheology and spontaneous fluctuations with fit of power laws. As a control measurement, we show that in the absence of both microtubules and actin, the active energy collapses. Concentration of cytochalasin B and nocodazole in interphase: 10 mg/ml. **d)** Analysis of the prefactor e0 of the power laws for the active energy show a significant decrease during pro- and metaphase as compared to interphase. Fit parameters to median shown in c) are displayed as blue dots. **e)** In contrast, the exponent *ν* of the fitted power laws remains constant throughout mitosis. Fit parameters to the medians as presented in c) are displayed as blue dots. n_*cells*_ = 58, 15, 19, 21, 20.

As the stiffness is reduced during metaphase, we hypothesize that the energy required for moving the probe particles is greatly reduced during mitosis. Indeed, comparing the prefactors of the power laws directly confirms this, as the cellular contribution to mobility decreases towards metaphase (Fig. 2d). After metaphase, the active energy used for particle mobility increases again and in telophase has almost returned to interphase levels. In contrast, the exponent of the power law seems unchanged throughout mitosis, suggesting that the processes driving particle motion remain unchanged, and only their amplitude is controlled by the cell (Fig. 2e). Potentially, intracellular mobility is reduced to a minimum to focus the cell’s energy on the division of the chromosomes or to reduce fluctuations that might interfere with the microtubules targeting the kinetochores. Alternatively, the reduced active mobility could just be a side effect of cytoskeleton remodeling that simply prevents significant transport during mitosis.

### Mechanics and transport switch from microtubule to actin during mitosis

Next we investigated the underlying structural reason for both the fluidification and the reduced active energy during mitosis. Therefore, we asked how actin and microtubules contribute to the mechanics during interphase and mitosis. To study this question, we decided to focus on cells that are in a mitotic arrest triggered by STC, a drug that keeps cells in mitosis after nuclear envelope breakdown and chromatin condensation. Hence, STC-treated cells structurally resemble mitotic cells in prophase. Furthermore, none of the six parameters that describe the mechanics and activity show a significant difference between STC-treated and prophase cells (Supplementary fig. S2). This confirms that we have a relevant model system for mitosis with respect to our measurements.

We systematically compared the effect of actin and microtubule depolymerizing drugs on the six passive and active mechanical parameters obtained in interphase and mitotic cells. Interestingly, inhibiting actin nucleation with cytochalasin B alone did not show any significant effect on the mechanics nor the active transport in interphase cells (Fig. 3). In contrast, treatment with nocodazole, which inhibits microtubule nucleation, significantly softened the cytoplasm and increased its fluid-like character as experienced by the probe particles (Fig. 3a-d). These results are further confirmed by simultaneous inhibition of actin and microtubule nucleation, which led to similar mechanical properties as when disrupting the microtubules alone. This supports the idea that actin does not play a crucial role for intracellular mechanics in interphase cells as experienced by membrane wrapped organelles. Similar results were obtained regarding the active motion driven by the active energy (Fig. 3e,f). While inhibiting actin nucleation did not lower intracellular activity, this activity was significantly reduced after inhibiting microtubule nucleation, and was fully abolished when both the actin and microtubules were depolymerized together (see also Fig. 2c). This is not a surprise, as basically no motor protein activity is possible when both actin and microtubules are absent.

**Figure 3:**
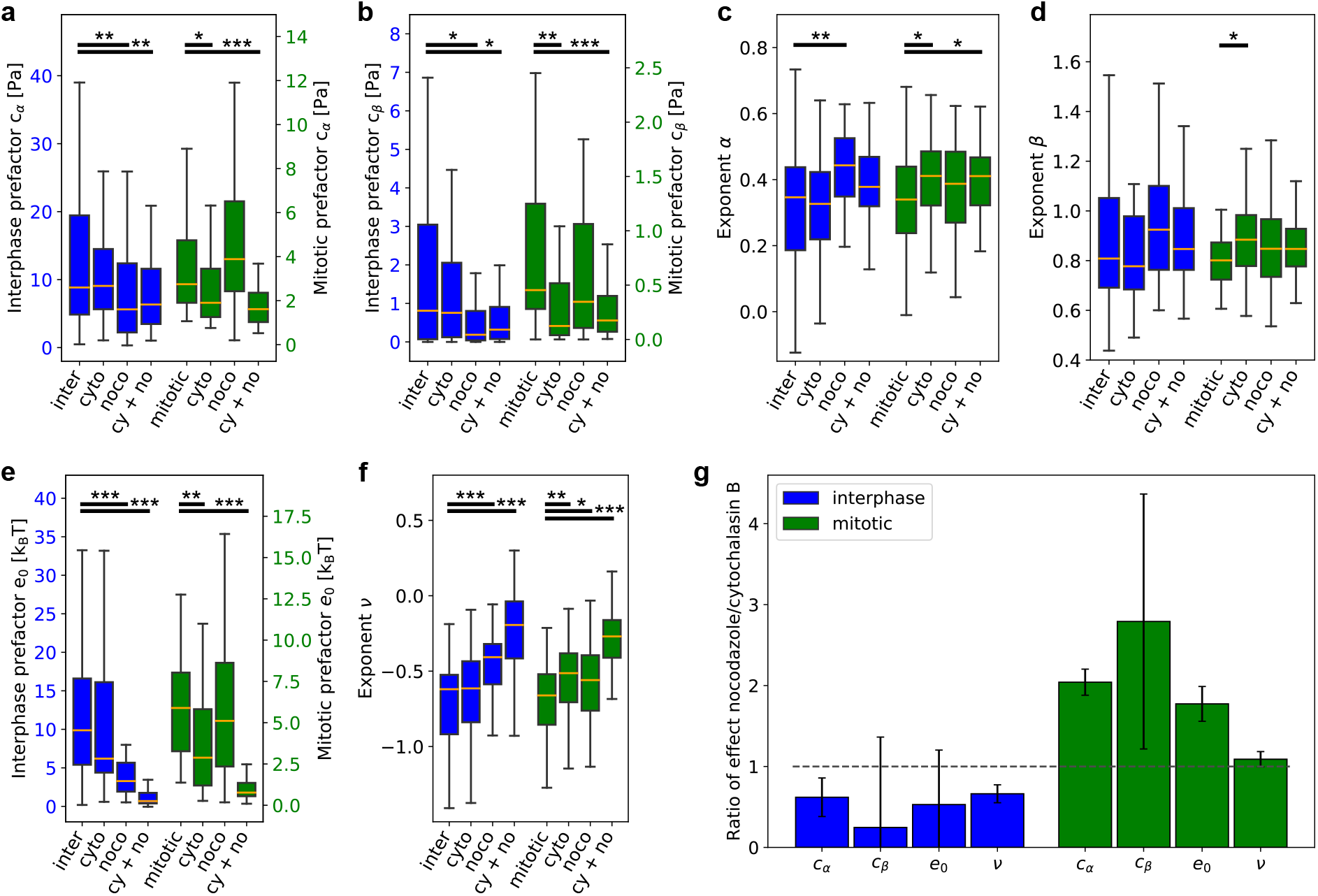
Comparison of viscoelasticity and activity in interphase and mitotic cells under cytoskeletal drugs. Influence of actin-interfering cytochalasin B and microtubule-interfering nocodazole (both at 10 μg/ml) on interphase and STC-treated cells (2 μM). **a)** The mechanical prefactor *c_α_* as an indicator for stiffness in low-frequency regime. Left y-axis (blue) for interphase cells, right y-axis (green) for mitotic cells (also for b and e). **b)** Mechanical prefactor *c_β_* as an indicator for stiffness in high-frequency regime. **c)** Power law exponent *α* as a measure of structure in low-frequency regime. **d)** Power law exponent *β* as a measure of structure in high-frequency regime. **e)** Prefactors *e*_0_ of power law fits to active energy as an indicator of intracellular activity. **f)** Exponents *ν* of power law fits to active energy as an indicator of intracellular activity. **g)** Ratio of nocodazole-treated cells to cytochalasin B-treated cells. A value less than 1 indicates a higher resulting parameter after nocodazole treatment, values higher than 1 indicate a higher parameter after cytochalasin B treatment. In interphase, nocodazole reduces intracellular stiffness and activity while in mitosis only cytochalasin B leads to this effect. Outliers not shown. *n_cells_* = 57, 43, 52, 61, 36, 55, 64, 66.

We then repeated these experiments in mitotic cells. To our surprise, we observed an opposite effect. Here, depolymerizing the actin network by cytochalasin B had a similar drastic effect on the stiffness as microtubule inhibition did in interphase cells (Fig. 3a,b). Besides the influence on stiffness, we also found fluidification to a comparable extent as well as a decrease of activity upon cytochalasin B-treated mitotic cells (Fig. 3d-f). However, inhibition of microtubule nucleation did not lead to a significant change of stiffness or activity. Furthermore, simultaneously inhibiting actin and microtubule nucleation led to a softening and fluidification comparable to actin inhibition alone. Similar to the intracellular activity in interphase cells, active transport was completely abolished in mitotic cells after combined actin and microtubule inhibition.

Our data suggests that the remodeling of the cytoskeleton during mitosis leads to a switch of mechanical roles between microtubules and actin with respect to organelle transport (Fig. 3g). While actin plays little to no role for intracellular mechanics during interphase, its contribution to the mechanical behaviour of the cell becomes dominant during mitosis. The profound changes after cytochalasin B treatment can be seen in all four parameters that describe the mechanical behaviour of the cell. The effect is relevant over the full frequency regime covered by our measurements. In addition, the reduction of mobility shows that transport processes that primarily acted via microtubules in interphase, switched to actin during mitosis. Again, this is surprising as the restructuring on the cytoplasm also seems to modify the motor proteins bound to the probe particles’ membrane wrapping. As such, we can speculate about possible mechanisms that effectively protect microtubules from transport vesicles during mitosis, potentially to ensure that the segregation of the chromosomes is not perturbed by transport of organelles. However, such protection would have to be complemented by an additional attachment mechanism to the sparse actin network in the cytosol to prevent their reorientation towards the cell cortex; this was not found within the scope of this study.

### Perturbation of actin leads to changes in intracellular mechanics and activity in dividing cells

The observed selective binding of the probe particles to the actin network during mitosis suggests that the structure of the cytosolic actin contributes to such organelle confinement roles, as the particles do not bind or redistribute to the actin-rich cortex. To further examine how actin nucleation or the motor protein myosin are responsible for the observed mechanical changes during mitosis, mitotic cells were treated with cytoskeletal drugs that block active force generation via myosin II (blebbistatin) and actin nucleation either via formins (SMIFH2) or the Arp2/3 complex (CK-666).

Blebbistatin shows a slight but not significant reduction of c_*α*_, indicating a stiffness decrease and a slight fluidification on long timescales, while the high-frequency regime as tested by c_*β*_ remains unchanged (Fig. 4a). Further, blebbistatin has drastic effects on the mobility, which is decreased to a similar level as after actin depolymerization. This demonstrates that the active motion as measured during mitosis is driven by myosin II, and not due to the ongoing cytoskeletal rearrangements inside dividing cells.

**Figure 4:**
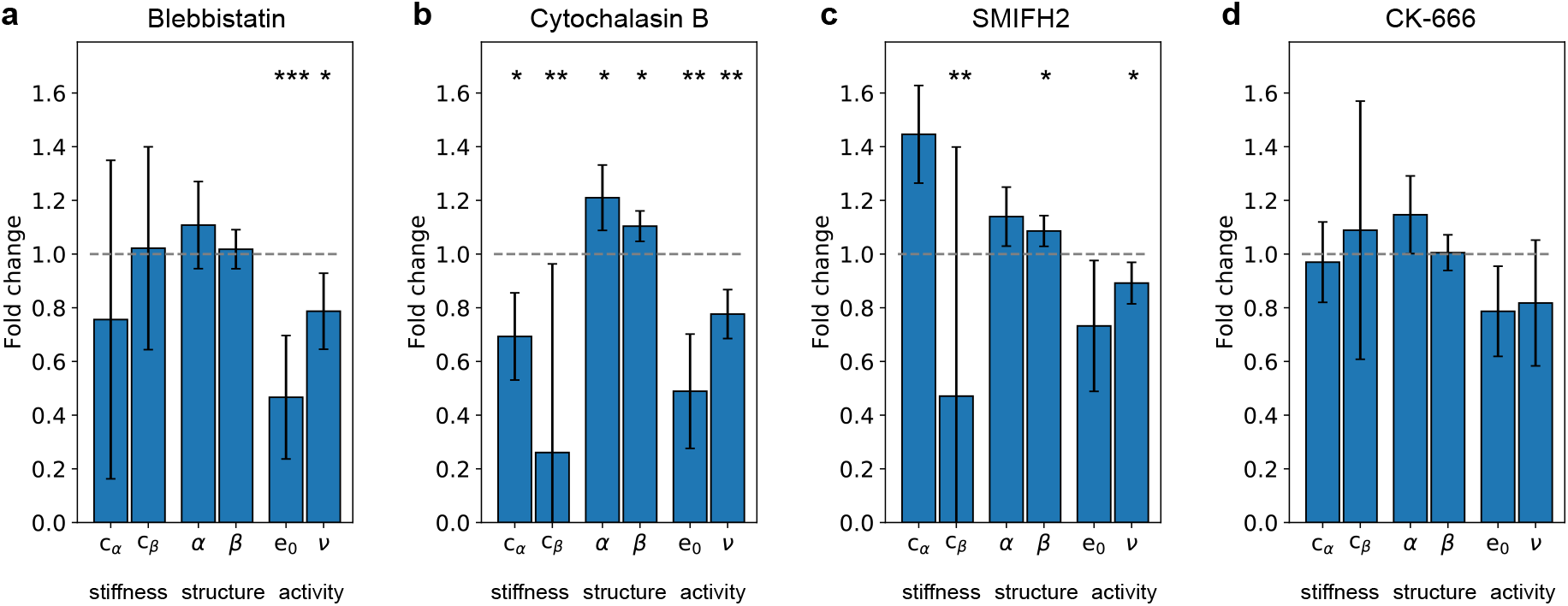
Intracellular mechanics and activity under actin perturbation. Actin drugs reveal great influence of formin-dependent and -independent actin nucleation in mitotic cells. Fold change of median compared to mitotic cells arrested by 2 μM STC after treatment with different inhibitors. **a)** Myosin II-inhibitor blebbistatin (20 μM), **b)** Actin nucleation inhibitor cytochalasin B (20.85 μMl), **c)** Formin inhibitor SMIFH2 (30 μM), **d)** Arp2/3 inhibitor CK-666 (100 μM). *n_cells_*: 34, 55, 54, 61.

Experiments in meiosis show that the intracellular, cytosolic network is largely nucleated by formins [37] while the cortical network is generated by the branching nucleator Arp2/3. Therefore, we wondered if a similar separation can be observed in mitosis. As Arp2/3-nucleated networks are highly branched and hence dense and stiff, our finding of decreased stiffness suggests that formin and not Arp2/3 is the main protein responsible for the cytosolic mechanics. Indeed, treatment with the formin inhibitor SMIFH2 shows an effect on the mechanical properties, while the inhibition of Arp2/3 did not significantly change the mechanics (Fig. 4c,d). A surprising result is that the inhibition of formins did influence the parameter c_*β*_ for the fluid component much more than c_*α*_. Although this tendency reflects our observation using cytochalasin B (Fig. 4b), it remains an open question as to why formin has such a precise effect on the fluid component of the viscoelastic cytosol. A possible explanation is that formin nucleated networks are not inherently cross-linked like Arp2/3 networks; hence, classical theories for entangled and not cross-linked networks may apply better to such a material.

### A new paradigm for intracellular mechanics during mitosis

Taken together, these results suggest a new view that cells actively switch both their mechanical and active intracellular processes during cell division (Fig. 5). While both the stiffness and active transport in interphase are comparably high, the restructuring during mitosis massively changes these processes. In stark contrast to the cortical stiffening and force increase, we observe a strong softening and fluidification of the cytoplasm that is accompanied by a reduction in the overall active transport of the probe particle during mitosis. Our experimental evidence shows that in interphase the mechanical properties as experienced by our probe particles are dominated by microtubules; however, during mitosis this switches to actin as the dominating component. This cannot be explained by a simple reduction of microtubules, as these are of course still present in mitosis, but take on the role of segregating the chromosomes. From a biological perspective, it makes perfect sense to protect the microtubules in this phase from organelles or membrane-engulfed particles. We suggest that during division, cells precisely control the interaction between organelles and actin or microtubules, leading to a targeting of our probe particle to the formin dependent f-actin. Then, the reduction of actin nucleation or myosin activity leads to substantial softening of the cytosol and to a decrease in intracellular active energy. As a consequence of these intracellular mechanical changes, the increased cortex stiffening can be interpreted as a protection mechanism that allows for a simplified way to segregate chromosomes while keeping the cellular integrity.

**Figure 5:**
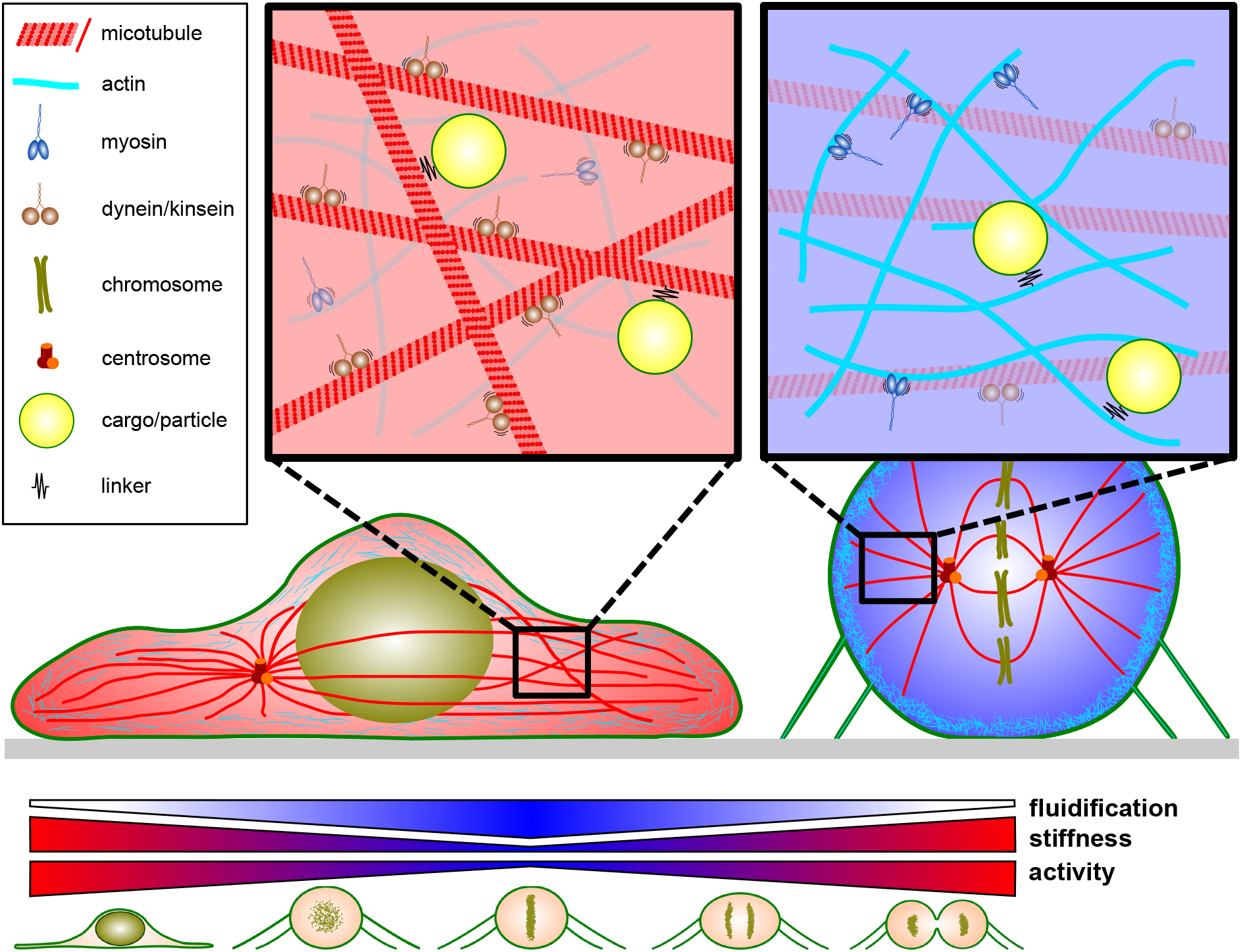
Model of intracellular mechanics in interphase and mitosis. In interphase, microtubules dominate the mechanics and are important for the intracellular activity. This switches to actin in mitosis, during which microtubules no longer substantially contribute to the intracellular mechanics or activity.

## Conclusion

Our experiments extend the knowledge about the mechanical properties in dividing cells from the cortex to the cytoplasm. By combining active optical tweezers-based microrheology with measurements of spontaneous particle motion, we gain full access to both the passive and active mechanical properties inside the cell. Using a two-material approach, we were able to quantify the full cellular mechanics over a large timescale with only six parameters. The model assumption of two independent, viscous and solid materials and a single active process predicts that these parameters can be changed independently of each other when affecting the respective relevant physical structures. Indeed, our experiments show that we can independently affect the relative contributions in the low- and high-frequency regime using different cytoskeletal drugs. In addition, during mitosis, exclusively the contribution of the solid-like, low-frequency component is reduced. Surprisingly, the solid-like material and the active energy depend not on a single physical element during the cell cycle: In interphase both are dominated by microtubules, and in mitosis this switches to actin (Fig. 5). This implies that both actin and microtubules can serve as structural support with the characteristics of a low-frequency, solid-like material. Furthermore, the intracellular activity is mostly microtubule driven in interphase, while it is actin driven in mitosis. The switch between microtubule to actin dominance could be explained by the vast remodeling of the cytoskeleton at the onset of mitosis.

The well-known cortical stiffening during mitosis was shown to be important for providing space for cell division, as highly confined cells show severe damage during mitosis [38]. However, another hypothesis for the relevance of cortical stiffening is that it serves as mechanical protection for the chromosomes and the spindle. Our observed softening and fluidification of the cytoplasm directly show that such a protection is indeed important, as otherwise the cell would have no internal mechanical support to resist external forces. Furthermore, the softening and fluidification suggests a physiological role of these changes, as it will greatly facilitate the mechanical processes during mitosis; future studies could test such a hypothesis by modifying the intracellular mechanical properties without disturbing the other, cell division-relevant processes.

## Materials and Methods

### MDCK and HeLa cell culture

In this study we used MDCK cells stably transfected with Lifeact-GFP and H2B-mCherry and HeLa cells transfected with H2B-mCherry (both cell lines kindly provided by Prof. R. Wedlich-Söldner). All cells were cultured in high glucose DMEM (Capricorn) supplemented with 10 % fetal bovine serum (Sigma-Aldrich), 1 % penicillin-streptomycin solution (Gibco) and 10 mM HEPES (AppliChem). Cells were kept at 37 °C and 5% CO_2_. To maintain selection pressure, transfected cells were kept in presence of 1 μg/ml puromycin.

Cells were seeded at low density (~ 10 000 cells/ml) on 24 × 50 mm coverslips with thickness of #1.5 (VWR). After a settling time to allow cells to adhere, ~ 50 μg/ml of polystyrene microparticles were added until the start of the experiment, but for at least 60 min. Excess particles that were not phagocytosed were washed off with prewarmed medium prior to the experiment. The coverslip was transferred onto a spacer slide with a thickness of 400 μm. Medium, optionally supplied with drugs, was added and the spacer slide was sealed with a 24 × 40 mm coverslip from the top. The slide was then transferred to the microscope and the setup was adjusted.

### Drugs and particles

All drugs were purchased from Sigma-Aldrich and were used at the indicated concentrations. Drugs were added 10 min prior to the experiment, except for STC, which was added at least 8 h prior to the experiment.

Polystyrene microparticles with a diameter of 1 μm, fluorescently labelled with DY-648P1 (653/672 nm) and NH_2_ modified, were purchased from Micromod (micromod Partikeltechnologie).

### Optical tweezers experiments

The optical tweezers used a near-infrared fiber laser (λ = 1064 nm, YLR-5-LP; IPG Photonics) that was position-controlled by a pair of acousto-optic XY-deflectors (DTSX-400-1064; AA Opto-Electronic) which were controlled by a variable frequency driver (Voltage Controlled Oscillator, DRFA10Y2X-D41k-34-50.110; AA Opto-Electronic). After deflection, the beam size was expanded to overfill the objective back aperture and directed into an inverted microscope (Eclipse Ti-e; Nikon Corporation) from the rear port. The beam was coupled in the optical path of the microscope by a dichroic mirror and focused in the object plane through a water immersion objective (Plan Apo VC WI, 60x, NA = 1.2; Nikon Corporation). To measure the force, the condenser was replaced by a force sensor module (Lunam T-40i; Impetux Optics), which was positioned according to the manufacturer procedure. The module was pre-calibrated and gives direct access to the force applied by the tweezers on any trapped object.

To achieve precise displacement measurements, a second, stationary near-infrared laser (λ = 976 nm, BL976-PAG500, Thorlabs) was used at low power (~ 0.25 mW). To determine the particle position, the detection laser is passed through the Lunam force detector, and the back focal plane of the condenser is imaged on a position sensitive detector (PDP90A, Thorlabs). The analog voltage signals of both the force detector and the position sensitive detector are acquired using an analog to digital acquisition card (BNC2090A, National Instruments Corporation) and further processed. To determine the particle position, the detection laser signal is calibrated using a stage scan over the particle via a piezo element (PXY 80 D12, piezosystem Jena) embedded in the motorized microscope stage (MicroStage Series, Mad city labs) before the measurements. A characteristic slope was fit to the scan to convert the voltage signal into micrometer displacement. This was done for every measured particle. All hardware was controlled using custom written LabVIEW programs (National Instruments Corporation).

### Microscopy and image acquisition

The optical tweezers were coupled to an inverted microscope (Eclipse Ti-e; Nikon Corporation), equipped with a custom built heating chamber, keeping it at 37 °C. The microscope was equipped with epifluorescence, as well as a spinning disk system (CSU-W1 Yokogawa; Intelligent Imaging Innovations Inc.) and a CMOS camera (Orca-flash4.0v2; Hamamatsu Photonics K.K.) for fluorescent imaging and a CCD camera (acA1920-155um, Basler AG) for brightfield imaging.

The DNA was recorded via the fluorescently labeled histones (H2B-mCherry), and the resulting images were used to categorize the different mitotic phases, prophase (condensed chromosomes), metaphase (chromosomes aligned in metaphase plate), anaphase (chromosomes separated) and telophase (cleavage furrow visible from actin signal/brightfield image).

### Active microrheology and cell mechanics

To perform active microrheology, phagocytosed micron-sized particle were trapped with the optical tweezers using a laser power of ~ 30 mW at the sample plane. The trap was then oscillated with a defined driving frequency and an amplitude of 200 nm. One measurement sequence contained a frequency sweep from 0.1 to 1000 Hz. The force applied to the particle was measured with the force sensor module and the stationary detection laser was used to measure particle displacement via laser interferometry with nanometer precision. From the phase shift between applied force and particle displacement, the viscoelastic response function *χ*\*(*f*) was calculated as:

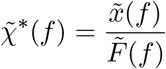

where *x* represents the measured particle displacement and *F* the applied force. 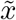 and 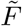 stand for the Fourier transform of the respective variable. During data analysis, data points that did not exceed the noise level by one standard deviation of data in the region one decade around the measured frequency were not taken into account. If more than two frequencies of a frequency sweep were rejected, the full measurement was discarded.

According to the generalized Stokes-Einstein theorem [19], the complex shear modulus *G*\* was calculated as:

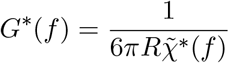

with *R* as the radius of the used micro-particle.

To fit the complex shear modulus, a fractional viscoelastic model with two power law components was used. This simplifies analysis and allows comparison between different phases and drug treatments:

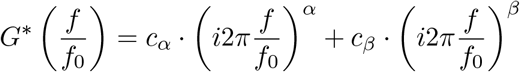

where *f*_0_ is the frequency of 1 Hz. Real and imaginary part of the complex shear modulus were simultaneously fit with G’ and G” as:

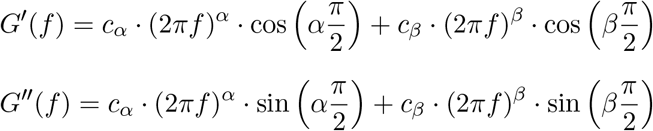

### Free particle fluctuations and cell activity

To determine the mobility of the particles without an optical trap, only the low power (~ 0.25 mW), stationary laser was used to measure the position of the particle over several 10 s time intervals. The positional fluctuations of the particle were converted to a power spectral density (PSD, 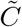) by:

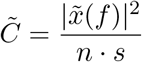

with *n* as the total number of data points and the sampling rate *s*. If the PSD reached a plateau at frequencies below 1000 Hz, the signal to noise ratio was not sufficient and the measurement was discarded.

To compare the measured response function (active microrheology) with the spontaneous particle fluctuations, a predicted response function assuming thermodynamic equilibrium can be calculated using the fluctuation dissipation theorem (FDT):

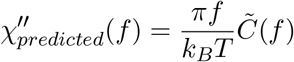

As the FDT is only valid in equilibrium, comparing the predicted with the determined response function allows one to dissect the extend of active forces on the particle motion. For the cells observed here, we find differences between the measured and the predicted response function in the lower frequencies (< 500 Hz), as expected for living active systems. The discrepancy in the low-frequency regime was termed active energy (*E_a_*), which was defined as:

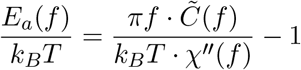

For higher frequencies, the cell reaches thermodynamic equilibrium, which means that responses calculated from spontaneous particle fluctuations and active microrheology overlap. As this was not the case for all measurements, mainly because of drifts in z-direction during spontaneous fluctuation measurements, the response calculated from passive measurements was shifted on top of the response calculated from active microrheology.

To simplify analysis and comparison between phases and drug treatments, a power law was fit to the active energy:

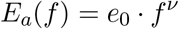

For the fit, frequencies with high noise levels for the particle fluctuations (around ~ 100 Hz) were not taken into account. The prefactor *e_0_* and exponent *ν* of the power law fit were used to compare the activity of cells for different phases and drug treatments.

### Data presentation and statistical analysis

Boxplots extend from first to third quartile, with the median value as center line. The whiskers extend from the edges of the box to the farthest data point in the 1.5 * interquartile range. Outliers are shown as dots, if not stated otherwise.

Median values are given with ± s.e.m. Statistical analyses were made using the two-sided t-test for parameters c_*α*_, e_0_ and *ν* (log normal distribution assumed and tested with D’Agostino-Pearson test for normality) and two-sided Mann-Whitney *U* -test for c_*β*_, *α* and *β* with *p*-values ≥ 0.05 not being considered significant. Significant *P* values were categorized into values * *p* < 0.05, ** *p* < 0.01 and *** *p* < 0.001. All measurements were repeated in at least three independent samples (N ≥ 3).

## Acknowledgements

We thank M. Brandt and T. Münker for their technical support and helfpul discussions. We are thankul for critical comments by Prof. E. Raz and M. Reichman-Fried. We thank Prof. R. Wedlich-Söldner for the MDCK II cell line expressing H2B-mCherry and GFP-Lifeact and HeLa line expressing H2B-mCherry. This work was supported by the European Research Council ERC-Consolidator grant PolarizeMe (771201).

## Author Contributions

S.H. carried out experiments and analyzed data. B.V. contributed to the experiments and with helpful discussions. T.B. designed and supervised the study. All authors wrote the manuscript.

## Data Availability

The data that support the findings of this study are available from the corresponding author upon reasonable request.

## Code Availability

The code that was used to analyse the data is available from the corresponding author upon reasonable request.

