## Supplementary Material for "Intracellular softening and fluidification reveals a mechanical switch of cytoskeletal material contributions during division"

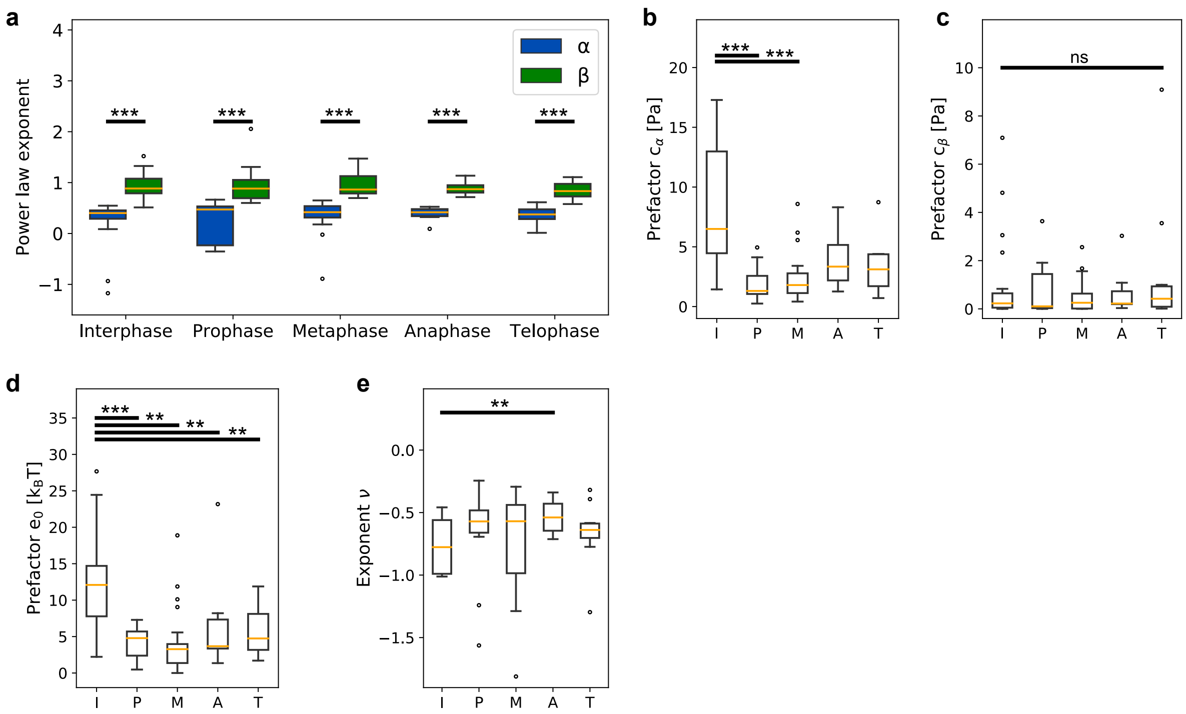


Figure S1: **Intracellular viscoelasticity and activity in dividing HeLa cells.** The parameters show a similar behaviour of mechanical properties as in MDCK cells. **a)** Power law exponents $\alpha$ and $\beta$ of viscoelastic fit for different phases of mitosis. **b)** Prefactor $c_\alpha$ of viscoelastic fit. **c)** Prefactor $c_\beta$ of viscoelastic fit. **d)** Prefactor $e_0$ of power law fit to active energy. **e)** Power law exponent $\nu$ of fit to active energy. $n_{cells}$: 20, 11, 23, 11, 10.

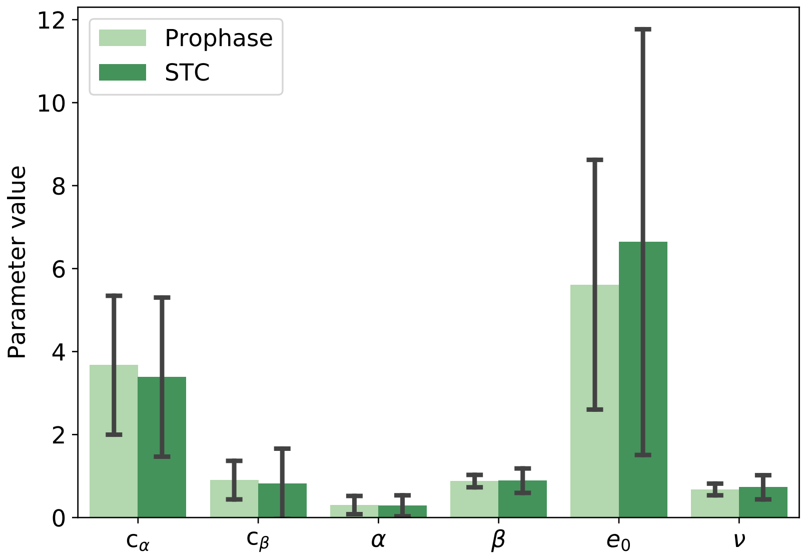


Figure S2: **Comparison of mechanical and active mechanical parameters of MDCK cells in prophase and arrested in mitosis by STC.** The parameters do not show a significant difference between cells in prophase and treated with STC, arresting the cells in mitosis. Data shows mean ± s.e.m.

| Phase | $c_\alpha$ [Pa] | $c_\beta$ [Pa] | $\alpha$ | $\beta$ | $e_0$ [$k_B$T] | $\nu$ |
|---|---|---|---|---|---|---|
| MDCK | | | | | | |
| Interphase | 8.84 ± 1.75 | 0.81 ± 0.41 | 0.35 ± 0.03 | 0.81 ± 0.04 | 9.88 ± 1.54 | -0.62 ± 0.04 |
| Prophase | 3.75 ± 0.43 | 0.92 ± 0.12 | 0.38 ± 0.06 | 0.84 ± 0.04 | 5.18 ± 0.78 | -0.69 ± 0.04 |
| Metaphase | 2.64 ± 2.26 | 1.31 ± 0.41 | 0.29 ± 0.08 | 0.77 ± 0.04 | 4.30 ± 0.60 | -0.65 ± 0.04 |
| Anaphase | 6.77 ± 1.46 | 1.27 ± 0.30 | 0.39 ± 0.04 | 0.91 ± 0.05 | 6.45 ± 0.84 | -0.65 ± 0.05 |
| Telophase | 8.69 ± 2.38 | 1.49 ± 0.61 | 0.34 ± 0.15 | 0.82 ± 0.04 | 7.75 ± 2.04 | -0.59 ± 0.06 |
| HeLa | | | | | | |
| Interphase | 6.49 ± 1.10 | 0.23 ± 0.41 | 0.40 ± 0.10 | 0.88 ± 0.06 | 12.09 ± 1.42 | -0.78 ± 0.05 |
| Prophase | 1.31 ± 0.43 | 0.10 ± 0.34 | 0.47 ± 0.12 | 0.88 ± 0.12 | 4.79 ± 0.70 | -0.57 ± 0.11 |
| Metaphase | 1.80 ± 0.40 | 0.25 ± 0.14 | 0.42 ± 0.06 | 0.86 ± 0.05 | 3.27 ± 0.91 | -0.57 ± 0.16 |
| Anaphase | 3.34 ± 2.52 | 0.22 ± 0.25 | 0.42 ± 0.04 | 0.87 ± 0.04 | 3.68 ± 1.72 | -0.54 ± 0.04 |
| Telophase | 3.11 ± 3.18 | 0.42 ± 0.86 | 0.38 ± 0.05 | 0.83 ± 0.05 | 4.74 ± 0.99 | -0.64 ± 0.08 |

Table S1: **Parameters for the two-material viscoelastic fits.** Median values for MDCK and HeLa cells for different phases of the cell cycle. Parameters of fit to $G^*(f) = c_\alpha \cdot (i2\pi f)^\alpha + c_\beta \cdot (i2\pi f)^\beta$ for mechanical parameters and $E_a(f) = e_0 f^\nu$ for the active energy. Errors represent s.e.m.
